# Factorial state-space modelling for kinetic clustering and lineage inference

**DOI:** 10.1101/2023.08.21.554135

**Authors:** R. Gupta, M. Claassen

## Abstract

Single-cell RNA sequencing (scRNAseq) protocols measure the abundance of expressed transcripts for single cells. Gene expression profiles of cells (cell-states) represent the functional properties of the cell and are used to cluster cell-states that have a common functional identity (cell-type). Standard clustering methods for scRNAseq data perform *hard* clustering based on KNN graphs. This approach implicitly assumes that variation among cell-states within a cluster does not correspond to changes in functional properties. Differentiation is a directed process of transitions between cell-types via gradual changes in cell-states over the course of the process. We propose a latent state-space Markov model that utilises cell-state transitions derived from RNA velocity to model differentiation as a sequence of latent state transitions and to perform *soft* kinetic clustering of cell-states that accommodates the transitional nature of cells in a differentiation process. We applied this model to the differentiation of Radial-glia cells into mature neurons and demonstrate the utility of our method in discriminating between functional and transitional cell-states.

## Introduction

Differentiation processes are typically represented as hierarchical transitions between functional cell-types. In contrast, it is widely assumed that scRNAseq data capture incremental shifts of gene expression along the differentiation process. Gene expression profiles of single cells measured using scRNAseq contain information about the functional properties of cell-states, and a differentiation model of incremental change of cell-states i.e. change in gene expression, is not consistent with the former model of discrete cell-type transitions. The latter model of differentiation allows for the existence of transitional cell-states that may be intermediate in both expression and function [1].

In standard scRNAseq analysis workflow, a k-nearest neighbours (KNN) graph of cell-states is used to cluster cells using community detection algorithms like Louvain or Leiden [2][3][4][5]. Clusters of cell-states are identified as cell-types based on marker gene expression and differential expression analysis. Grouping cells in this manner aids interpretability; however, since cell-states within a cluster are all assigned the same label, variation within the clusters is implicitly discarded as uninteresting noise in further analyses.

For scRNAseq data from differentiation processes, lineage inference models seek to impose a transitional relationship between cell-states. The models compute pseudotime, a score representing progress along the differentiation axis and also partition cell-states between multiple co-occurring lineages. Pseudotime estimation can model incremental shifts in gene expression, but due to the high sparsity of scRNAseq data, inferring the functional properties of individual cell-states from gene expression is challenging. Therefore, the clustering of cell-states remains important for interpretation.

Prior work has attempted to develop models of cell-type transitions, mainly by building minimum spanning trees among cell-state clusters [6][7] or by aggregating the connectivity between cell-states for each cluster [8]. The underlying clustering itself is not informed by cell-state transitions since a KNN graph is undirected and symmetric; in contrast, the directed signal obtained from RNA velocity enables the estimation of transition probabilities between cell-states. This information can be represented as a directed and asymmetric graph [9].

We introduce a latent state-space model based on cell-state transitions that enable the clustering of cells based on their transition dynamics. We refer to this form of clustering as kinetic clustering. Cell state transitions are assumed to be observed emissions from dynamics in a smaller latent state-space. The model allows for the probabilistic *soft* assignment of cells towards latent states and can be used to identify transitional cell-states. The dynamics of the differentiation process are captured by transitions between latent states. Multiple lineages can be further modelled with additional factors representing parallel sequences of latent state transitions [10].

## Method

### Model Input

RNA velocity of single cells can be used to estimate transition probabilities among cell-states [9][11]. For the set of measured cell-states (observed states) *O* = {*o*^(1)^, …, *o*^(*n*)^ }and an initial probability vector *Y*_0_ = {*P* (*o*^(1)^ *i* = 0), …, *P* (*o*^(*n*)^ *i* = 0) }, the transition probability matrix **T** over observed states is used to simulate the differentiation process as a sequence of probability vectors **Y**. The simulation is performed as,

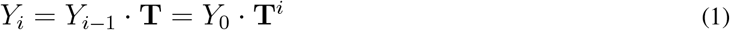

The simulation is considered to have converged if *Y*_*i*_ = *Y*_*i*_*−*_1_, i.e. when the simulation reaches the stationary state of the Markov chain with the transition matrix **T**. The latent dynamic model considers the simulated process,

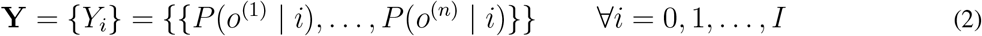

as input. In the following text, *P*_*o*_ is used to indicate a probability vector over states *O* such as *Y*_*i*_ = *P*_*o*_(*o* | *i*).

### Model Specification

With latent states *S* = {*s*^(1)^, …, *s*^(*m*)^ }and analogous to the simulation over observed states, we describe the dynamics over latent states as

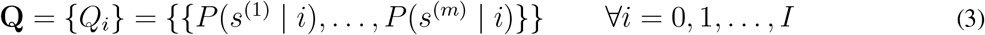

Let **H** be the transition probability matrix over latent states *S*, then corresponding to the simulation (Eq. (1)), a Markov chain in the latent space has the form,

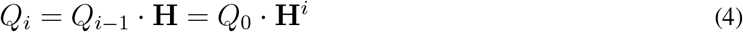

With the assumption of constant emission probabilities of observed states over the latent process *P* (*o* | *s, i*) = *P* (*o* | *s*), we express *Y*_*i*_ as

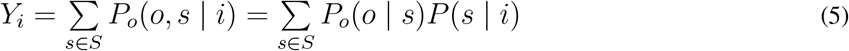

and due to Eq. (4):

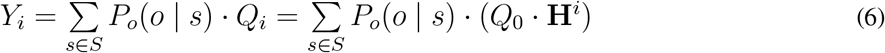

Lineages *L* are modelled as independent Markov chains in the latent space. Furthermore, restricting the lineages to a common latent state-space *P* (*o* | *s, l*) = *P* (*o* | *s*),

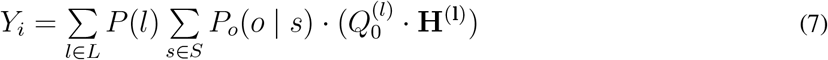

where **H**^(**l**)^ is the latent state transition probability matrix for lineage *l* ϵ *L* and,

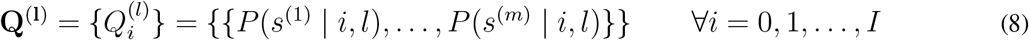

### Model Training

The trainable parameters of the model are the conditional latent state transition probability matrices **H**,

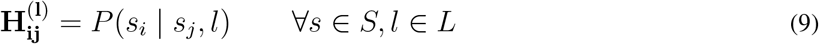

the emission probabilities **E**,

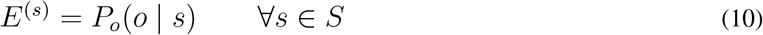

the lineage weights *W*,

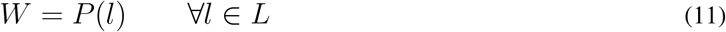

and the initial latent state probabilities **Q**_**0**_,

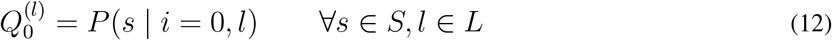

Let Ŷ be the model estimate of **Y**. The estimated sequence Ŷs obtained as,

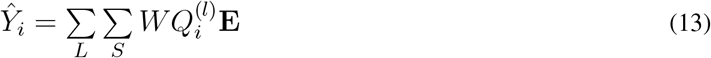

The parameters of the model are optimized by minimising the element-wise Kullback–Leibler (KL) divergence using gradient descent.

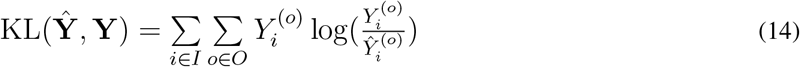

The training is regularised for sparsity in the latent state transition matrix with the addition of element-wise KL divergence between the diagonal value of the latent transition matrix and a vector of ones to the loss.

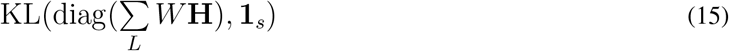

In order to obtain non-redundant latent states, in lieu of model selection, the Jensen-Shannon divergence of the conditional probability vectors of observed states given latent states is minimised.

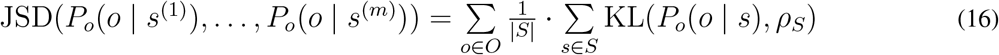

where,

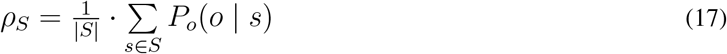

### Model Output

The pseudotime of any cell *o* ϵ*O* is estimated as the mean step of a cell weighted by the probability of observing a cell at each step *i*,

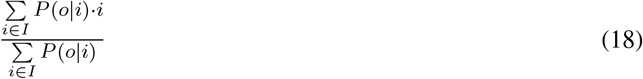

The conditional probability of latent states with respect to observed states (cells) is used to assign kinetic cluster memberships,

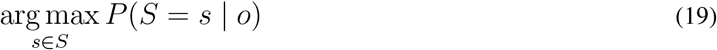

The transition entropy of a cell is the sum of the entropy of the joint probability of a cell and each latent state,

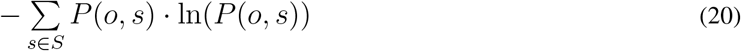

Each lineage *l* ϵ *L* is a sequence of transitions in a common state space *S*. The trajectories of lineages in latent state space are represented as sequences of most probable latent states at each step *i*,

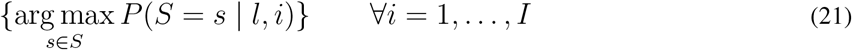

### Data availability

### Developing human forebrain

Data was downloaded from *scvelo v0*.*2*.*5*

### Code availability Notebooks

All datasets were processed using standard scRNAseq and RNA velocity workflow. Details of the analyses can be found

in the following notebooks. https://github.com/aron0093/cy2path_notebooks.

### Implementation

The model was implemented using PyTorch and Python code for the project can be found at the following repository https://github.com/aron0093/cy2path.

## Results

### Factorial state space modelling for differentiation processes

The observed state-space is considered to be discrete and composed of all observed cell-states. The differentiation process is conceptualized as the evolving probability distribution over the observed states and is modelled by simulating the RNA velocity-derived transition probability matrix of cell-states. The purpose of the latent state-space model is to create an interpretable summary of the simulation.

Under the latent state-space model, the differentiation process is the transitions between discrete latent states with probabilistic emissions of observed states. The model is parameterised with a transition probability matrix over latent states and emission probabilities of observed states for each latent state. Analogous to the simulation over observed states, the dynamics over latent states are learnt by minimizing the divergence between the simulation and the estimate from the latent state-space model.

Dynamics over the latent states are more interpretable since the number of latent states is much lower than the observed states. The probabilistic assignment of cells towards latent states is referred to as kinetic clustering. Kinetic clustering of cells is based on state transitions unlike clustering based solely on gene expression profiles. Kinetic clusters group cell-states that arise together during the differentiation process. Lineages are modelled as transitions between latent states and are also informative of the relative persistence of latent states. Multiple co-occurring lineages are modelled as independent Markov chains in latent state-space and can be considered independent components of the observed differentiation dynamics.

### Identifying transitional cells in developing human forebrain

The developing human forebrain dataset consists of the glutamatergic neuronal lineage in human embryonic cells. The process follows a linear differentiation path from Radial-glia (progenitor) cells via a neuroblast (intermediate) population that is locked into the neuron (mature) fate [9]. The intermediate neuroblasts are highly motile cells that migrate to target brain regions before terminal differentiation [12] [13].

Root and terminal states inferred with RNA velocity correspond to the Radial-glia and mature neurons, respectively, as has been previously reported [Figure 1A.3-4] [9]. The model was fit using default parameters [Notebooks]. Kinetic clustering partitioned the data into two clusters [Figure 1B.1]. Static clusters computed using the Leiden algorithm overlap exclusively with one of the kinetic clusters except Leiden cluster 3, which appears to be split between the two kinetic clusters [Figure 1B.3].

**Figure 1:**
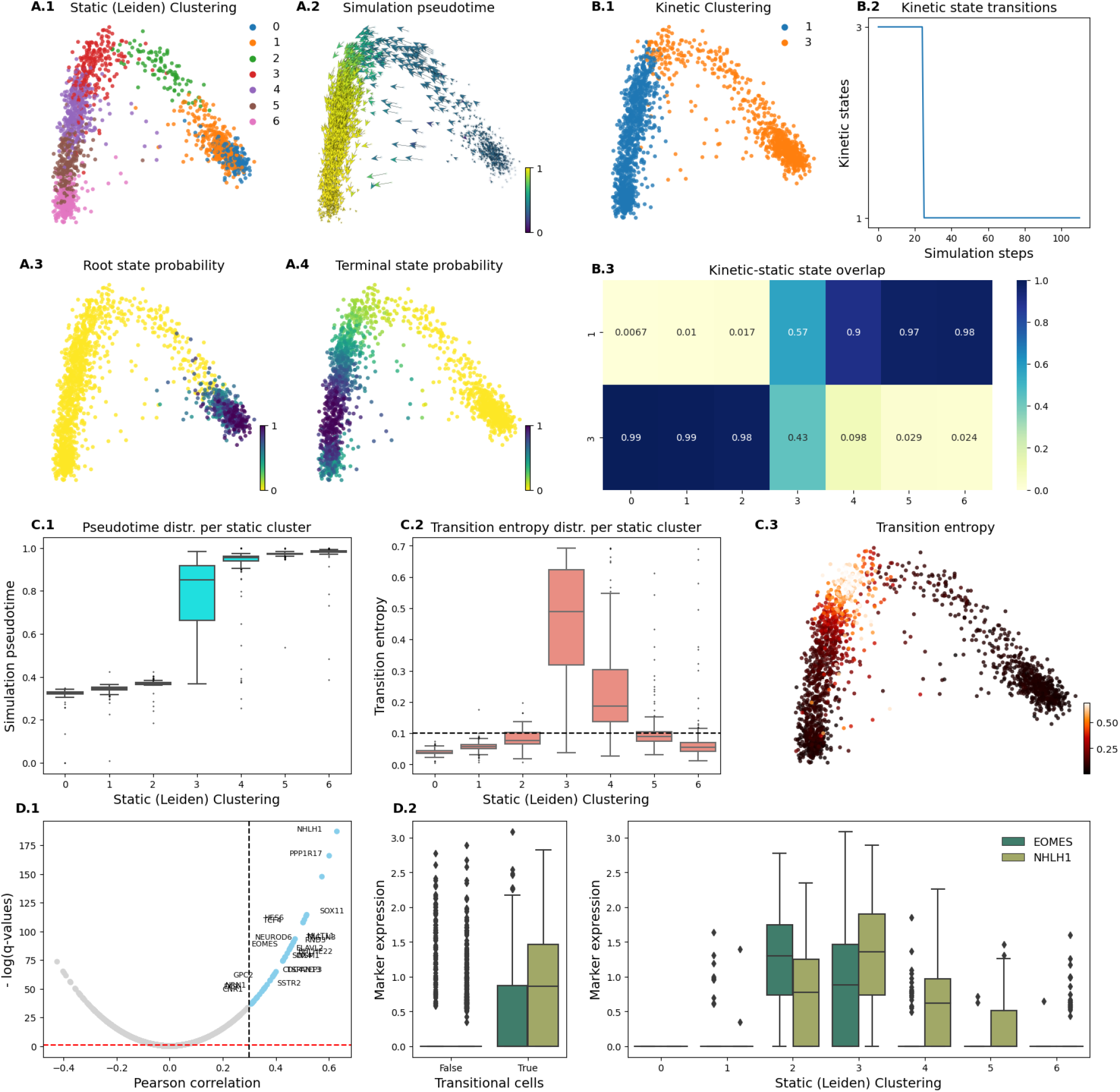
Identifying transitional cells in developing human forebrain. **(A)** Outputs of standard workflow scRNAseq and RNA velocity analysis annotated on the first two principal components. (A.1.) Leiden clustering of cell states, (A.2.) RNA velocity vectors and estimated pseudotime. The pseudotime of a cell is calculated as the mean step weighted by the probability of observing a cell at each step. (A.3.) Root and (A.4.) Terminal cell states inferred using RNA velocity. **(B)** Outputs of our latent state space model. (B.1.) Kinetic clustering of cell states which is the most probable latent state per cell. (B.2.) Most probable latent state at each simulation step. (B.3.) Ratio of overlapping cells in each static (Leiden) and kinetic cluster. **(C)** Identification of transitional cell states. (C.1.) Pseudotime distribution of cells in each static cluster. (C.2.) Transition entropy of cells; computed as the entropy of the joint probability of a cell and each latent state; distribution over static clusters. The red line is the threshold to discriminate transitional cells from the rest. (C.3.) Transition entropy of cells. **(D)** Biological identification of transitional cells as neuroblasts. (D.1.) Pearson correlation between gene expression and transitional entropy of cells. (D.2.) Marker genes’ (*EOMES, NHLH1*) expression distribution in transitional cells vs rest and for each Leiden cluster.

Pseudotime estimated using RNA velocity has high variance in Leiden cluster 3, suggesting that this cluster may contain transitional cells [Figure 1C.1]. Transitional cells were identified as cells with high transitional entropy [Figure 1C.2-3]. Marker genes for neuroblasts were enriched in the set of genes positively correlated with transitional entropy [Figure 1D.1] [14].

Cells high in the expression of *EOMES* [9] and of *NHLH1* [15], canonical markers for neuroblast cells, are spread across multiple Leiden clusters. Cells expressing marker genes overlap with cells identified as transitional [Figure 1D.2]. This analysis concludes that transitional entropy is a useful criterion for selecting transitional cells.

## Discussion

Differentiation processes are generally represented as a sequence of transitions between cell-types. Intermediate cell-types represent distinct, physiological stages of the process. Clustering of cell-states based on gene expression profiles is an essential step in the study of both terminally differentiated and differentiating cells. Groups of cells obtained in this manner are identified as canonical cell-types by marker identification and functional analysis. In contrast, lineage inference approaches utilise the highly resolved measurement of cell-states with scRNAseq, to model cell-state transitions as gradual processes and not as discrete transitions between cell clusters [16][11]. These methods infer differentiation coordinates for individual cells in the form of pseudotime and cell fate probability.

Inference of the identity and function of individual cell-states is challenging due to dropout of genes measured in scRNAseq and high biological stochasticity between similar cell-states. Dimensionality reduction is a necessary step to compensate for missing measurements by exploiting correlation in genes’ expression. While dimensionality reduction reduces the number of features, clustering is an analogous process of reducing the number of states. Cells within a cluster are assigned the same label and subsequent analysis such as differentiation expression testing implicity assumes variation between these cell-states to be uninformative variation. The reduction of states aids interpretabiltiy and discovery of functional associations between genes.

For scRNAseq data from differentiation processes, unlike terminally differentiated cells, some variation within clusters corresponds to the differentiation process itself. A model of transitions between clusters, while interpretable, cannot faithfully represent an incremental process as well as gradual divergence of lineages. Information on transitional cells, the relative time of transitions and the persistence of intermediate states is lost in such a representation [8][6][7].

The asynchronous differentiation of cells is the fundamental basis for constructing models of differentiation processes from scRNAseq data. It can therefore be expected that biological samples collected at different time points will have different distributions of cell-states. Therefore, we propose an approach that models the differentiation process as an evolving probability distribution over observed cell-states. Such an approach allows for the simultaneous persistence of cell-states in different stages of the process while also capturing the sequence of transitions as well as the relative temporal coordinate. RNA velocity of single cells has enabled the estimation of asymmetric transitions between cell-states and several methods have used these transitions for lineage inference [9][11]. In prior work, we demonstrated the utility of a Markov simulation-based lineage inference approach that exploits emergent properties not discernable via analytical formulations [17]. While simulations over cell-states have the desired properties for our modelling approach to differentiation processes, a simulation over several thousand cell-states is not interpretable.

Therefore, we introduce a latent state-space model where we consider the simulation over cell-states to be driven by a latent Markov process over a much smaller number of latent states. The inferred latent process has higher interpretability while retaining the attributes of our approach to differentiation processes. Kinetic clustering of cells is based on state transitions, unlike clustering based on only expression profiles. In the analysis of the human forebrain dataset, we demonstrate that kinetic clustering groups cells with temporal similarity and the utility of this approach in the identification of transitional cells. The model has been implemented using pyTorch and can make use of GPU parallelisation. The code relies on standard packages in the field and can easily be incorporated into RNA velocity-based trajectory inference workflows.

